# Temporary cerebral ischaemia impairs thromboxane A2 constriction and induces hypertrophic remodelling in peripheral mesenteric arteries of hypertensive rats: limited reversal despite long-term suberoylanilide hydroxamic acid cerebroprotection

**DOI:** 10.1101/2024.10.15.618453

**Authors:** Andrea Díaz-Pérez, Silvia Lope-Piedrafita, Belén Pérez, Paula Vázquez-Sufuentes, Maria Rodriguez-Garcia, Ana M Briones, Xavier Navarro, Clara Penas, Francesc Jiménez-Altayó

**Author notes:** **Correspondence:** Dr. Francesc Jiménez-Altayó, Departament de Farmacologia, de Terapèutica i de Toxicologia, Facultat de Medicina, Institut de Neurociències, Av. de Can Domènech, s/n, Universitat Autònoma de Barcelona, 08193, Bellaterra (Barcelona), Spain.

## Abstract

Stroke induces brain injury, especially severe in hypertensive patients, and elevates mortality rates through non-neurological complications. However, the potential effects of a transient ischaemic episode on the peripheral vasculature of hypertensive individuals remain unclear. Here, we investigated whether transient cerebral ischaemia (90 min)/reperfusion (1 or 8 days) induces alterations in mesenteric resistance artery (MRA) properties in adult male spontaneously hypertensive rats (SHR). In addition, we assessed whether the reported cerebroprotective effects of suberoylanilide hydroxamic acid (SAHA; 50 mg/kg; administered intraperitoneally at 1, 4, or 6 h after reperfusion onset) extend long-term and include beneficial effects on MRAs. Functional and structural properties of MRAs were examined at 1- and 8-days post-stroke. Nuclei distribution, collagen content, and oxidative stress were assessed. Ischaemic brain damage was evaluated longitudinally using magnetic resonance imaging. Following stroke, MRAs from SHR exhibited non-reversible impaired contractile responses to the thromboxane A2 receptor agonist U46619. Stroke increased the MRA cross-sectional area, wall thickness, and wall/lumen ratio due to augmented collagen deposition. These changes were partially sustained 8 days later. SAHA did not improve U46619-induced contractions but mitigated stroke-induced oxidative stress and collagen deposition, preventing MRA remodelling at 24 h of reperfusion. Furthermore, SAHA induced sustained cerebroprotective effects over 8 days, including reduced brain infarct and oedema, and improved neurological scores. However, SAHA had minimal impact on chronic MRA contractile impairments and remodelling. These findings suggest that stroke causes MRA changes in hypertensive subjects. While SAHA treatment offers long-term protection against brain damage, it cannot fully restore MRA alterations.

## 1. Introduction

Stroke stands as one of the leading causes of death and adult disability globally [1, 2]. Hypertension is the principal modifiable risk factor for ischaemic stroke. Clinical and observational studies underscore that maintaining low blood pressure levels correlates with a reduced incidence of both initial and recurrent ischaemic strokes [3]. Stroke patients often experience non-neurological complications that can variably damage peripheral organ systems such as the lungs, kidneys, and gastrointestinal tract, which can hamper post-stroke brain recovery [4]. However, there is still a gap in understanding how the deprivation of blood to the brain affects peripheral tissues.

The combination of stroke and gastrointestinal complications significantly worsens morbidity and mortality rates in patients [5]. Mesenteric resistance arteries (MRAs) play a crucial role in supplying blood to the small intestine and most of the colon [6]. Alongside other small resistance arteries and arterioles, they constitute the peripheral vascular resistance, which significantly influences blood pressure regulation in conjunction with cardiac output [7]. Hypertension, associated with a surge in cardiovascular disease, is often linked to disturbances in the systems governing resistance artery tone. These disturbances cause endothelial dysfunction, excessive constriction, and vascular remodelling, leading to cardiovascular pathologies [8, 9]. Essential hypertension is linked to an increase in total peripheral resistance, primarily attributed to vascular remodelling [10, 11]. Small resistance arteries in patients with essential hypertension often exhibit inward eutrophic remodelling [12]. This process involves the reorganization of normal arterial material around a constricted lumen, leading to an increased media/wall-to-lumen ratio, which clinically predicts cardiovascular events independent of blood pressure [13]. Mesenteric resistance arteries from spontaneously hypertensive rats (SHRs) show a consistent increase in wall-to-lumen ratio and wall thickness, suggesting that this strain is a suitable model for studying hypertension-related structural alterations [14].

After a cerebral ischaemic event, blood supply can be partially or fully restored during a phase called reperfusion. This period is crucial as it aims to replenish oxygen and nutrients to the affected brain tissue. However, it can also trigger reperfusion injury, characterized by inflammation and oxidative damage due to the sudden return of blood flow [9]. Beyond the brain, reperfusion injury prompts the mobilization of neutrophils, T lymphocytes, and platelets, which adhere to the endothelium, as well as the release of reactive oxygen species (ROS) and pro-inflammatory mediators [15, 16]. Consequently, the systemic surge in oxidative and pro-inflammatory mediators induces redox stress, potentially affecting the non-ischaemic hemisphere, as well as organs and blood vessels distant from the brain [4, 17, 18]. Disruption in thromboxane A2 signalling is linked to stroke pathophysiology [19, 20], however, its specific impacts on the cerebral and peripheral vasculature remain uncertain. Stroke patients exhibit impaired peripheral vasodilation due to endothelium-dependent and endothelium-independent mechanisms [21, 22]. Notably, previous studies have linked experimental cerebral ischaemia (I)/reperfusion (R)-induced interleukin-6 production with increased ROS, leading to MRA endothelial dysfunction in normotensive rats [23]. Nevertheless, the precise impact of cerebral I/R injury on the peripheral vascular beds of hypertensive individuals, who already have underlying vascular changes, are not fully understood.

The histone deacetylase (HDAC) inhibitor suberoylanilide hydroxamic acid/Vorinostat (SAHA), approved by the U.S. Food and Drug Administration for treating cutaneous T-cell lymphoma, has shown cerebroprotective effects in murine ischaemic stroke models when used at doses equivalent to those for treating cancer [24, 25, 26]. Our recent study demonstrated that administering SAHA within 4 h of reperfusion onset offers cerebroprotection in SHRs after 24 h of reperfusion [27]. Interestingly, a previous study indicated that administering SAHA between 4 and 10 days after a focal photothrombotic stroke in normotensive mice led to sustained functional recovery by enhancing the neuroplasticity of surviving neurons in the peri-infarct zone through epigenetic mechanisms [25]. Yet, the potential long-term beneficial effects of SAHA administered during reperfusion on brain infarct and stroke-induced peripheral alterations remain unexplored, particularly in hypertensive animals.

In the present study, our aim was to investigate whether transient cerebral I (90 min) followed by R (1 or 8 days) induces changes in MRAs from SHR, and to determine if these changes are reversible. Furthermore, we sought to examine whether the cerebroprotective effects of SAHA extend to the long term and include beneficial effects on the peripheral vasculature.

## 2. Material and Methods

### 2.1. Animals

A total of 72 adult male SHR, aged 12-14 weeks, were used. Animals were housed under a 12 h-dark/light cycle, had free access to food and water, and were kept in controlled environmental conditions. All the experiments were carried out in accordance with the guidelines established by the Spanish legislation on protection of animals used for scientific purposes (RD 53/2013) and the European Union Directive (2010/63/UE). The protocols received approval from the ethics committee of the Universitat Autònoma de Barcelona (approval code: CEAAH 4275M4).

### 2.2. Experimental design

A first set of 55 rats was used to evaluate tMCAO-induced alterations in the functional and structural properties of MRAs in SHR subjected to 90 min of ischaemia followed by 24 h of reperfusion. The experimental design, exclusion criteria for surgical outcome, and the cerebroprotective effects of SAHA were reported in a previous study [27]. Briefly, rats were intraperitoneally (*i*.*p*.) injected at 1, 4, or 6 h from reperfusion onset with a single dose of SAHA (50 mg/kg; LC laboratories, Woburn, MA, USA, Cat#: V-8477) or vehicle composed of 10% DMSO (Sigma-Aldrich, San Luis, MO, USA, Cat#: D5879), 40% PEG-300 (Sigma-Aldrich, Cat#: 81162), 5% Tween-80 (Sigma-Aldrich, Cat#: P4780), and 45% saline (Alco, Terrassa, Spain, Cat#: 141659.1211). Animals were randomly allocated into the following groups: tMCAO + VEH (*n* = 7), tMCAO + SAHA 1 h (*n* = 5), tMCAO + SAHA 4 h (*n* = 5), and tMCAO + SAHA 6 h (*n* = 4). A sham-operated group was used (SHAM; *n* = 5), in which the same surgical procedure was performed, but nylon suture was removed 1 min after occlusion.

A second set of 17 rats was used for the longitudinal assessment of the potential cerebroprotective effects of SAHA, as well as for the analysis of MRA wall properties in SHR subjected to 90 min of ischaemia followed by 8 days of reperfusion. Rats were administered *i*.*p*. with the same volume of vehicle (as above) or SAHA (50 mg/kg) 4 h after the onset of reperfusion. The time-point and dose of administration was established based on previous results [27], which showed clear cerebroprotection in the group injected with SAHA 4 h after reperfusion. The dosage also followed previous studies [24, 27]. Animals were randomly allocated to the different experimental groups using random number generator software (Graphpad Prism®, version 9.0; GraphPad Software Inc., San Diego, CA, USA). Investigators were blinded to the treatment assignment during and after surgical procedures. The following experimental groups were established: tMCAO + VEH (*n* = 7) and tMCAO + SAHA 4 h (*n* = 6). Four animals were excluded from the study according to the following criteria: technical surgical complications (*n* = 2) and brain haemorrhage (*n* = 2). Figure 1 provides an overview of the animal cohorts employed for both the long-term assessment of the brain effects of SAHA and the study of long-term MRA properties. To determine the appropriate sample size, a calculation was conducted using brain infarct data obtained from a prior study [27], which evaluated the acute cerebroprotective effects of SAHA. This calculation was performed using Institutional Animal Care and Use Committee software at the University of Illinois Urbana-Champaign, Chicago, IL, USA. Subsequently, it was determined that a sample size of 5–7 animals per group would be adequate to detect statistically significant differences, with a significance level (*p*) set at 0.05 and a statistical power of 80%.

**Figure 1.**
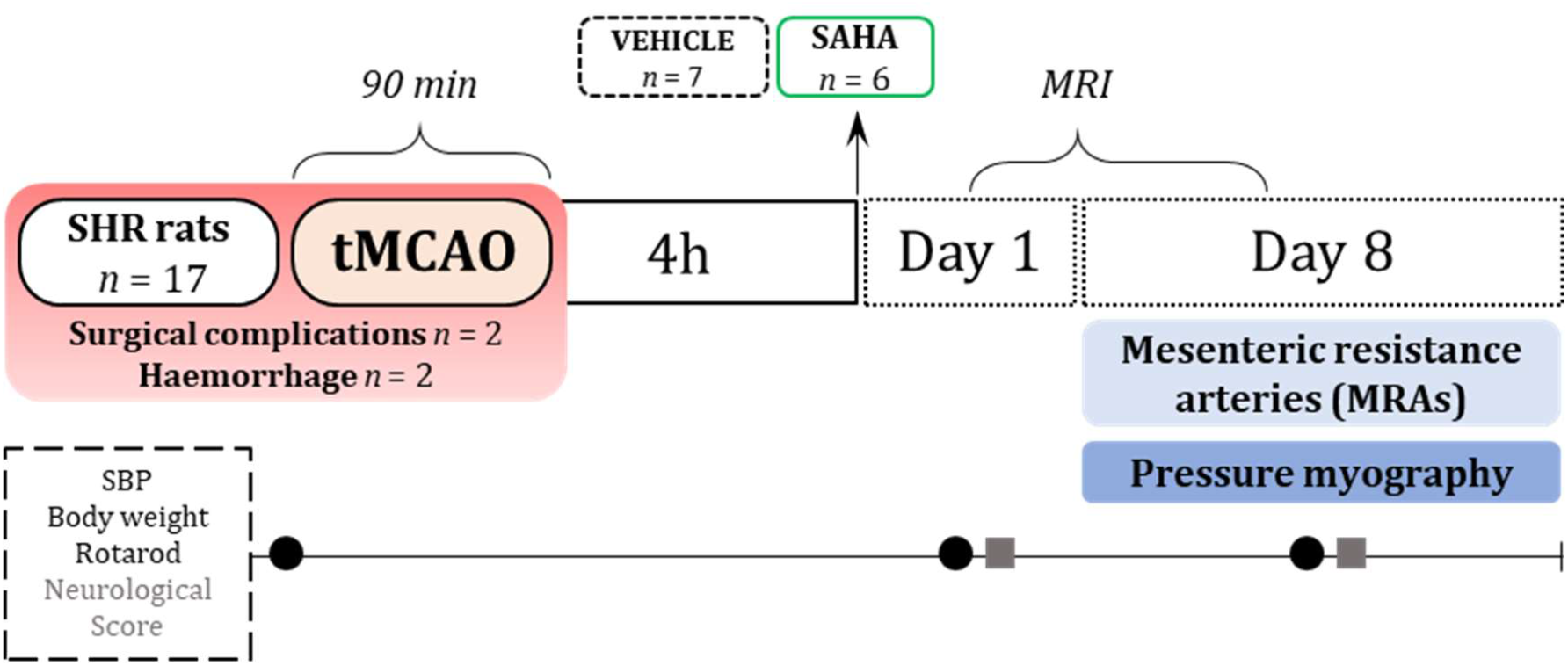
CONSORT-style flow diagram of the long-term assessment of the brain effects of SAHA and the study of long-term MRA properties, illustrating the animals used, the study design, and experiment timeline. Spontaneously hypertensive rats (SHR) underwent 90 min of transient middle cerebral artery occlusion (tMCAO) followed by 8 days of reperfusion. A single dose of suberoylanilide hydroxamic acid (SAHA; 50 mg/kg) or an equivalent volume of vehicle was administered 4 h after the onset of reperfusion. Four animals were excluded based on the following predefined criteria: i) technical surgical complications; and ii) brain haemorrhage due to mechanical damage. Ischaemic brain damage was evaluated longitudinally using magnetic resonance imaging (MRI) on days 1 and 8 of reperfusion. Systolic blood pressure (SBP), body weight, and performance in the rotarod test were assessed 1 day before surgery and 1 and 8 days after reperfusion. Neurological outcomes were evaluated on days 1 and 8 after reperfusion. After 8 days of reperfusion and following euthanasia, mesenteric resistance arteries were dissected free of fat and connective tissue and mounted in a pressure myograph to study their functional and structural properties.

### 2.3. Blood pressure measurements

Systolic blood pressure (SBP) was measured in conscious SHR using the tail-cuff method (NIPREM 645; Cibertec, Madrid, Spain), as reported [27]. Rats were placed in a dark, warm cage for at least 10 min, and a tail pulse sensor and pressure cuff were situated around their tails. When the pulse was detectable, the cuff was inflated and deflated several times, and finally, the SBP was recorded. Before the final measurements (1 day before surgery, and 1- and 8-days post-reperfusion), rats were habituated to the dark, warm cage for 3 days. Measurements were always performed within the same hour range. The average SBP of each rat was determined from 3 repeated measurements. Rats were considered hypertensive when SBP values where ≥ 160 mmHg.

### 2.4. Induction of transient middle cerebral artery occlusion (tMCAO)

The right MCA was occluded for 90 min using a nylon suture thickened at the end (RWD-MCAO, Biogen Científica, Madrid, Cat#: MSRC40B200PK50), as described [27, 28]. Anaesthesia was induced with 5% isoflurane (Zoetis Spain S.L.U., Madrid, Spain, Cat#: EX5085-19-01) vaporized in O_2_ and N_2_O (at a ratio of 30:70) and maintained at 2.5-3% isoflurane throughout the operation. In the case of sham-operated animals (control group), the same surgical protocol was followed, but the nylon suture was removed 1 min after occlusion. A thermal probe (heating blanket connected to a temperature-control system set at 37 ± 0.5°C; Cibertec, Madrid, Spain) was used to monitor the body temperature of the animal during the entire surgical procedure. Cortical cerebral blood flow (cCBF) was also monitored throughout the process using a laser Doppler flowmetry system (Perimed AB, Järfalla, Sweden) to corroborate MCA occlusion, as reported [29]. To monitor cCBF, a holder was implanted the day before the occlusion at specific coordinates relative to bregma (2 mm posterior and 3.5 mm lateral) from the thinned skull. At the end of the reperfusion period, 24 h or 8 days (Figure 1), animals were deeply anaesthetized with 5% isoflurane (vaporized in O_2_ and N_2_O at a ratio of 30:70) and euthanised.

### 2.5. Neurological outcomes

The 9 points-score test was conducted on day 1 and 8 after reperfusion to evaluate neurological outcomes (0 = no deficit, 9 = maximum impairment), as described [30]. In addition, the rotarod test (Harvard apparatus Ltd., Holliston, MA, USA) was employed to assess motor coordination and balance [31]. Briefly, animals were placed in the device at a fixed rate for 1 min before undergoing 3 cycles of acceleration (from 4 to 40 rpm), with resting intervals of 10 min between cycles. Each trial ended when the animal fell from the rotating cylinder, and the time spent prior to falling was recorded. Animals were habituated for 3 days before final measurements, which were taken 1 day before surgery and 1 and 8 days after occlusion. The average was calculated from three repeated measures each time point.

### 2.6. Tissue preparation

At 24 h after reperfusion, blood samples were collected via intracardiac puncture from deeply anesthetized animals (using 5% isoflurane vaporized in O_2_ and NO_2_ at a ratio of 30:70). The blood samples were centrifugated at 3000 rpm for 10 min, and plasma (500 μl) was collected, quickly frozen, and stored at −70°C for further analysis by high-performance liquid chromatography (HPLC) [27]. Following euthanasia, after 24 h or 8 days (Figure 1) of reperfusion, the mesentery was dissected and placed in cold Krebs-Henseleit (KH) solution (composition in mM: NaCl 112.0; KCl 4.7; CaCl_2_ 2.5; KH_2_PO_4_ 1.1; MgSO_4_ 1.2; NaHCO_3_ 25.0, and glucose 11.1). Third-order branches of the superior MA were dissected free of fat and connective tissue in ice-cold KH solution. One artery was immediately used for pressure myography and subsequently fixed in paraformaldehyde (Sigma-Aldrich, Cat#: 158127) for studies including nuclei distribution using microscopic fluorescence and collagen content analysis [32]. Another artery, which was not fixed in paraformaldehyde, was used for the evaluation of *in situ* oxidative stress [33].

### 2.7. Pressure myography

After 24 h or 8 days (Figure 1) of reperfusion, functional and structural properties of the MRA were evaluated using pressure myography (Danish Myo Technology, Aarhus, Denmark), as described [32]. Briefly, the vessels were mounted on 2 glass microcannulas and carefully positioned to ensure parallel vessel walls without stretching. Intraluminal pressure was increased to 140 mmHg, and the artery was unbuckled by adjusting the cannulas. Subsequently, the artery was allowed to equilibrate for 1 h at 70 mmHg in gassed KH solution at 37 °C. Intraluminal pressure was then reduced to 3 mmHg, and a pressure–diameter curve ranging from 3 to 120 mmHg was obtained. Internal and external diameters were measured for 3 min at each intraluminal pressure. To assess viability, the tissue was contracted with 100 mM KCl (Alco, Cat#: 191494.1211), followed by washing and a 30-min resting period. Constricted tone was induced using the stable thromboxane A2 mimetic 9,11-dideoxy-9α, 11α -methano-epoxy prostaglandin F2α (U46619; 1 nM–1 μM; EMD Millipore, Billerica, MA, USA, Cat#: 538944), before the addition of the endothelial-dependent relaxation agonist, acetylcholine (ACh; 1 nM– 100 μM; Sigma-Aldrich, Cat#: A6625). The artery was then equilibrated for 30 min at 70 mmHg in gassed, calcium-free KH solution at 37 °C (0 Ca^2+^: omitting calcium and adding 10 mM EGTA; Sigma-Aldrich, Cat#: E4378) (passive conditions), and a second pressure-diameter curve (3–120 mm Hg) was obtained. Finally, the artery was fixed (70 mmHg) with 4% of paraformaldehyde for 45 min and prepared for nuclei distribution analysis. Structural and myogenic parameters were obtained according to the calculations described previously [34].

### 2.8. Nuclei distribution

Pressured-fixed MRAs were stained with the nuclear dye Hoechst 33342 (0.01 mg/ml; Sigma-Aldrich, Cat#: 14533) for 30 min. After washing with phosphate-buffered saline, the artery was mounted on slides with silicone well spacers to prevent artery deformation [33]. The distribution of nuclei along the different arterial wall layers was visualized using a laser confocal Microscope (60x; FV1000; Olympus Iberia, Barcelona, Spain). Stacks of images were obtained, corresponding to 0.5 μm-thick serial slices from the adventitia to the lumen of the artery. At least two stacks of serial images were obtained from each artery and analysed using MetaMorph Image Analysis software (Molecular Devices). The shape and orientation of stained Hoechst 33342 cell nuclei allowed for differentiation between the different layers of the arterial wall. Wall volume, layer volumes (adventitia, media, and intima), and the number of cell nuclei present in each of the previously mentioned layers were determined, as described [34]

### 2.9. Collagen content

Pressure-fixed frozen MRAs from nuclei distribution studies were sliced into sections 14 μm thick and then incubated with Picrosirius Red solution: 0.1% Sirius Red/Direct Red 80 (Sigma-Aldrich, Cat#: 365548) in distilled water, 1.296% in picric acid (Sigma-Aldrich, Cat#: 74069) for 1 h to determine the total collagen content present in the arterial wall [32]. After incubation, any excess of dye was removed by rinsing the samples with acidified water (0.5% acetic acid in distilled water), and the samples were mounted for further observation. The total collagen content was quantified using Image J software (National Institutes of Health, Bethesda, MD, USA) by measuring the stained area in images obtained under visible light using an FV1000 optical microscope (Olympus Iberia). Measurements were taken from at least three rings from each animal, and the results were expressed as arbitrary units.

### 2.10. Oxidative stress measurements

The analysis of circulating 2-hydroxyethidium (2-EOH; Noxygen Science Transfer & Diagnostics GmbH, Germany, Cat#: NOX-14.1) in plasma was conducted using HPLC with fluorescence detection, serving as a quantitative measure of plasma superoxide anion levels, as previously described [31, 35, 36]. The presence of 2-EOH in the samples was quantified by comparison with a calibration curve based on the xanthine-xanthine oxidase reaction, following the method described previously [37].

*In situ* oxidative stress levels were assessed in 14-μm thick MRA cross sections using the oxidative fluorescence dye dihydroethidium (DHE; Sigma-Aldrich, Cat#: 37291) [37]. Briefly, DHE (2 μM) was applied onto each section, cover-slipped, and then incubated for 30 min in a humidified chamber at 37°C, protected from light. To ensure the specificity of the fluorescence signal, the antioxidant compound Mn(III)tetrakis(1-methyl-4-pyridyl)porphyrin (1.434 mg/ml; EMD Millipore, Cat#: 475872) was used as negative control. Images were captured using a laser confocal microscope (60x; FV1000, Olympus Iberia) with uniform acquisition settings across all experimental conditions. The average fluorescence intensity was quantified using ImageJ software in at least 2 sections from each artery.

### 2.11. Magnetic resonance imaging (MRI)

H-Magnetic resonance studies were conducted using a 7T Bruker BioSpec 70/30 USR system (Bruker BioSpin GmbH, Karlsruhe, Germany), equipped with a mini-imaging gradient set (400 mT/m), a linearly polarized transmit volume coil (72 mm inner diameter), and a dedicated rat brain circularly polarized surface coil as the receiver, as described [31]. MRI data acquisition and processing were performed on a Linux computer using Paravision 5.1 software (Bruker BioSpin GmbH). Briefly, rats were anaesthetized with 1.5% (v/v) isoflurane in O_2_ at 1 L/min and positioned in an animal bed with bite-bar and ear-bars for optimal head immobilization. Body temperature and respiration were continuous monitored using a small animal monitoring system (Model 1025; SA Instruments, Inc., Stony Brook, NY, USA), with body temperature maintained at 37°C using a circulating heated water bath [31].

For brain volume measurements, high resolution T_2_-weigthed (T_2_w) coronal images covering brain tissue from the olfactory bulb to the cerebellum were acquired using a Rapid Acquisition with Relaxation Enhancement (RARE) sequence with the following parameters: repetition time (TR) of 6500 ms ; echo spacing of 12 ms; RARE factor of 8; two echo images with effective echo times (TEeff) of 36 and 132 ms; four number of averages; 30 contiguous slices with a thickness of 0.5 mm and a 0.05 mm gap between slices; field of view (FOV) of 32 mm × 32 mm; and acquisition matrix (MTX) of 256 × 256. The scan duration was 10 min and 24 s. For volume estimations, image slices containing a lesion component were identified, and regions of interest were manually delineated to measure areas of cortical and subcortical lesions in the ipsilateral and contralateral hemispheres on each slice. Area measurements were multiplied by slice thickness plus the gap to calculate volume. Infarct volume was corrected for the effects of oedema by dividing the lesion volume by the ratio of ipsilateral to contralateral volume, as described [31].

### 2.12. Statistical analysis

The results are presented as mean ± SEM, except in Figure 7D, where they are expressed as median [Q1; Q3], with the number of animals (*n*) specified in each table/figure. Data distribution was consistently assessed before selecting the appropriate statistical tests. For analyses involving a single factor, one-way ANOVA followed by Bonferroni’s *post hoc* test, was employed for comparisons among more than two means. In cases involving two factors, a repeated or regular measures two-way ANOVA with Tukey’s *post hoc* test for grouped analyses was performed. The specific statistical tests used are indicated in each table/figure. Data analysis was conducted using GraphPad Prism® version 9.0 (GraphPad Software Inc., San Diego, CA, USA), with statistical significance considered at *p* < 0.05.

## 3. Results

### 3.1 Cerebral I/R impairs U46619-induced contractions in MRAs, unaffected by SAHA treatment

To evaluate whether cerebral I/R alters MRA properties, SHR were submitted to 90 min of MCA occlusion followed by 24 h of reperfusion, as previously reported [27]. A single dose of SAHA (50 mg/kg) or an equal volume of vehicle were administered (*i*.*p*.) at 1, 4, or 6 h after reperfusion onset. First, functional experiments were conducted to determine whether cerebral I/R impairs MRA relaxation and constriction. No significant differences in KCl (100 mM)-induced constrictions were observed among all experimental conditions (results not shown). The contractile responses of MRAs to increasing concentrations of the thromboxane A2 receptor agonist, U46619, were impaired after cerebral I/R, and this impairment was not prevented by SAHA treatment (Figure 2A). On the other hand, relaxation in response to ACh was similarly healthy regardless of I/R or SAHA treatment (Figure 2B).

**Figure 2.**
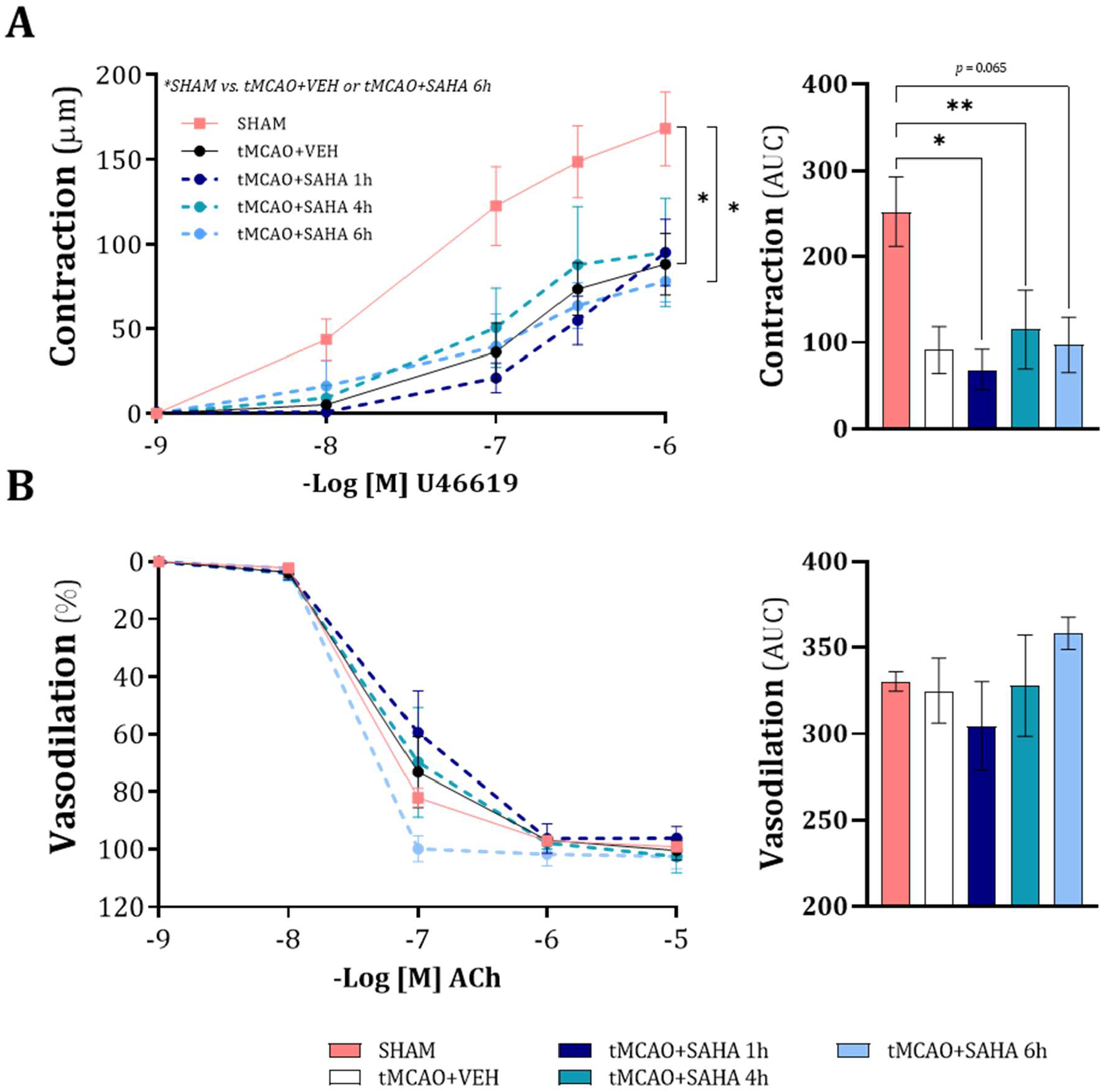
Influence of cerebral ischaemia (90 min)/reperfusion (24 h) (tMCAO) and SAHA treatment at 1, 4, or 6 h of reperfusion on MRA concentration-response curves to (A) U46619 and (B) acetylcholine (ACh) in hypertensive rats. The analysis of area under the curve (AUC) is presented on the right. Results are the mean ± SEM from SHAM (*n* = 5), tMCAO + VEH (*n* = 7), tMCAO + SAHA 1h (*n* = 5), tMCAO + SAHA 4h (*n* = 5), and tMCAO + SAHA 6h (*n* = 4). \**p* < 0.05; \*\**p* < 0.01 by repeated measures two-way ANOVA (left) or one-way ANOVA with Bonferroni’s *post hoc* test (right).

### 3.2. Cerebral I/R induces MRA remodelling: protective effects of SAHA treatment

MRA external and lumen diameters were measured across a pressure range of 3 to 120 mmHg under passive conditions (0 Ca^2+^-KH solution). We then assessed the impact of cerebral I/R and *in vivo* treatment with SAHA. Cerebral I/R did not significantly modify arterial diameters (Figure 3A and 3B). Administration of SAHA during reperfusion was associated with a reduction in both external (Figure 3A) and lumen (Figure 3B) diameters.

**Figure 3.**
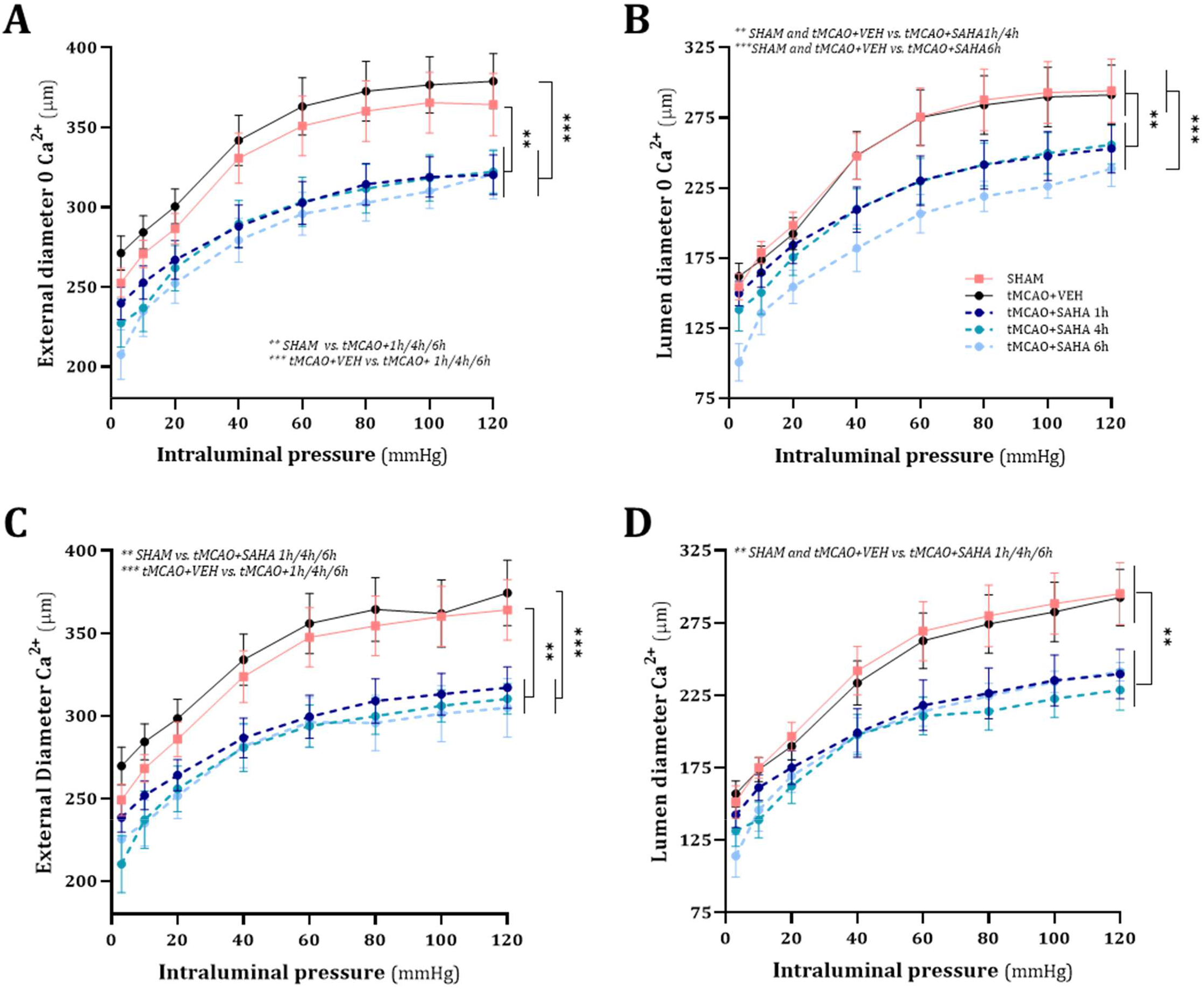
Influence of cerebral ischaemia (90 min)/reperfusion (24 h) (tMCAO) and SAHA treatment at 1, 4, or 6 h of reperfusion on MRA external and lumen diameters obtained in passive and active conditions (0 Ca^2+^ – Krebs-Henseleit solution) in hypertensive rats. External (A) and lumen (B) diameter – intraluminal pressure in passive conditions. External (C) and lumen (D) diameter – intraluminal pressure in active conditions. Results are the mean ± SEM from SHAM (*n* = 5), tMCAO + VEH (*n* = 7), tMCAO + SAHA 1h (*n* = 5), tMCAO + SAHA 4h (*n* = 5), and tMCAO + SAHA 6h (*n* = 4). \*\**p* < 0.01; \*\*\**p* < 0.001 by repeated measures two-way ANOVA.

Arterial diameters in active conditions (2.5 mM Ca^2+^-KH solution) remained unchanged by cerebral I/R (Figure 3C and 3D). On the contrary, vessel and lumen diameters decreased across the pressure range in SAHA-treated rats (Figure 3C and 3D). The myogenic response (i.e., internal diameter in active relative to passive conditions) of the MRA was nearly negligible and unaffected by cerebral I/R or treatment (results not shown).

Notably, under passive conditions, cerebral I/R augmented wall thickness (Figure 4A), cross sectional-area (Figure 4B), and wall-to-lumen ratio (Figure 4C) compared to sham operated SHR. Treatment with SAHA significantly decreased wall thickness when administered at 1 and 4 h post-reperfusion onset (Figure 4A), while it reduced cross-sectional area regardless of the timing of administration (Figure 4B), ultimately restoring them to levels comparable to those observed in sham-operated SHR. In addition, SAHA treatment attenuated the wall-to-lumen ratio increase when administered within 4 h of reperfusion onset (Figure 4C). Taken together, cerebral I/R induces hypertrophy in MRAs of SHR, an effect that is prevented by SAHA treatment, particularly when administered within 4 h of the onset of reperfusion.

**Figure 4.**
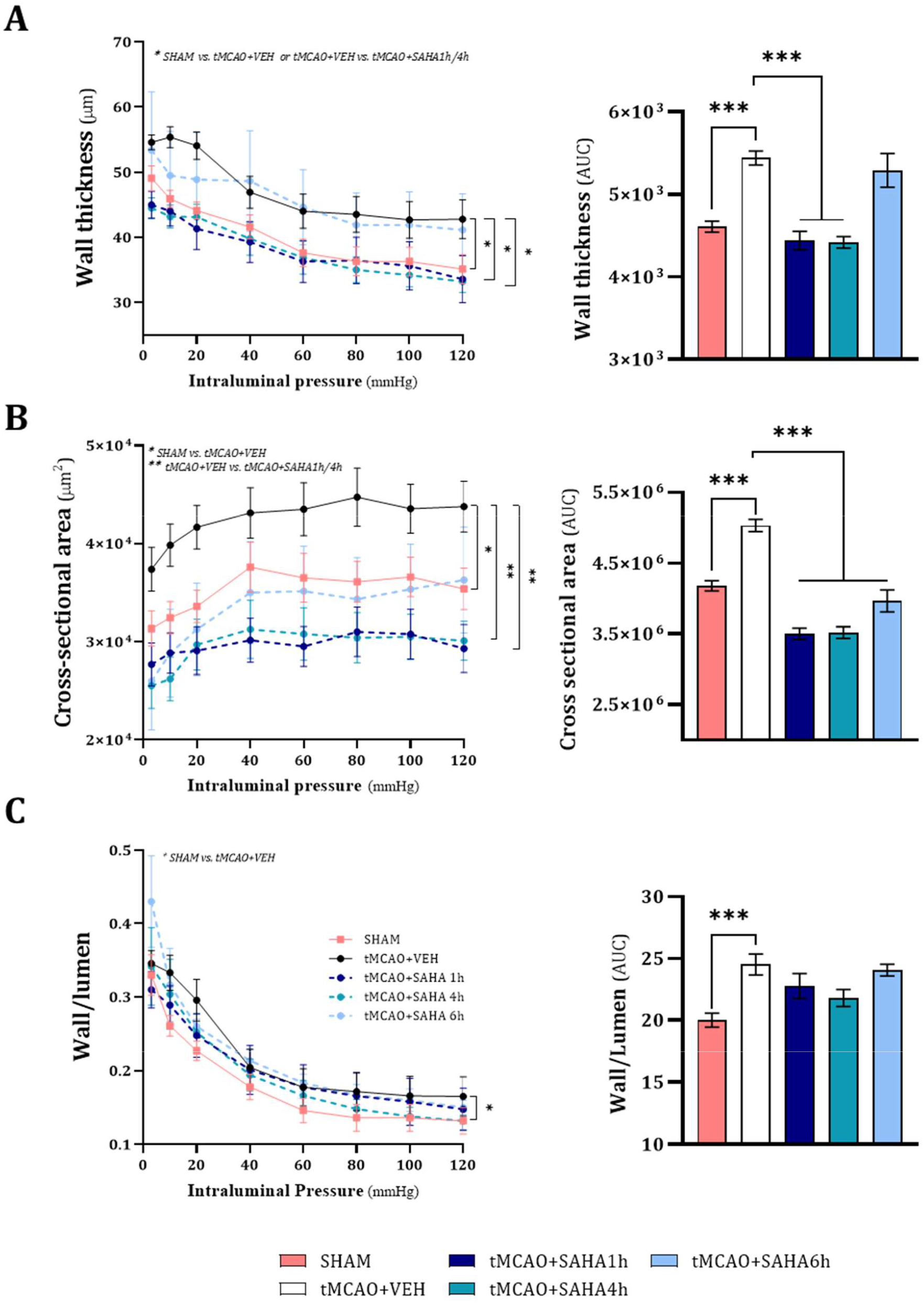
Influence of cerebral ischaemia (90 min)/reperfusion (24 h) (tMCAO) and SAHA treatment at 1, 4, or 6 h of reperfusion on MRA structural properties obtained in passive conditions (0 Ca^2+^ – Krebs-Henseleit solution) in hypertensive rats. (A) Wall thickness – intraluminal pressure. (B) Cross-sectional area – intraluminal pressure. (C) Wall/lumen ratio – intraluminal pressure. The analysis of area under the curve (AUC) is presented on the right. Results are the mean ± SEM from SHAM (*n* = 5), tMCAO + VEH (*n* = 7), tMCAO + SAHA 1h (*n* = 5), tMCAO + SAHA 4h (*n* = 5), and tMCAO + SAHA 6h (*n* = 4). \**p* < 0.05; \*\**p* < 0.01; \*\*\**p* < 0.001 by repeated measures two-way ANOVA (left) or one-way ANOVA with Bonferroni’s *post hoc* test (right).

### 3.3. Cerebral I/R leads to an increase in MRA media volume and collagen deposition, which is prevented by SAHA treatment

To explore the mechanisms underlying cerebral I/R-induced MRA wall hypertrophy and the protective effects of SAHA, we examined the distribution of nuclei across various wall layers using confocal microscopy in pressurized MRAs (Figure 5A). Quantitative measurements revealed that cerebral I/R increased wall, adventitial, and media volumes, but only the increase in media volume reached statistical significance (Figure 5B). Cerebral I/R did not significantly modify the number of adventitial cells, whereas a higher count of SMCs was observed without reaching statistical significance (Figure 5B). Notably, SAHA treatment reduced adventitial and media volumes and the number of SMCs.

**Figure 5.**
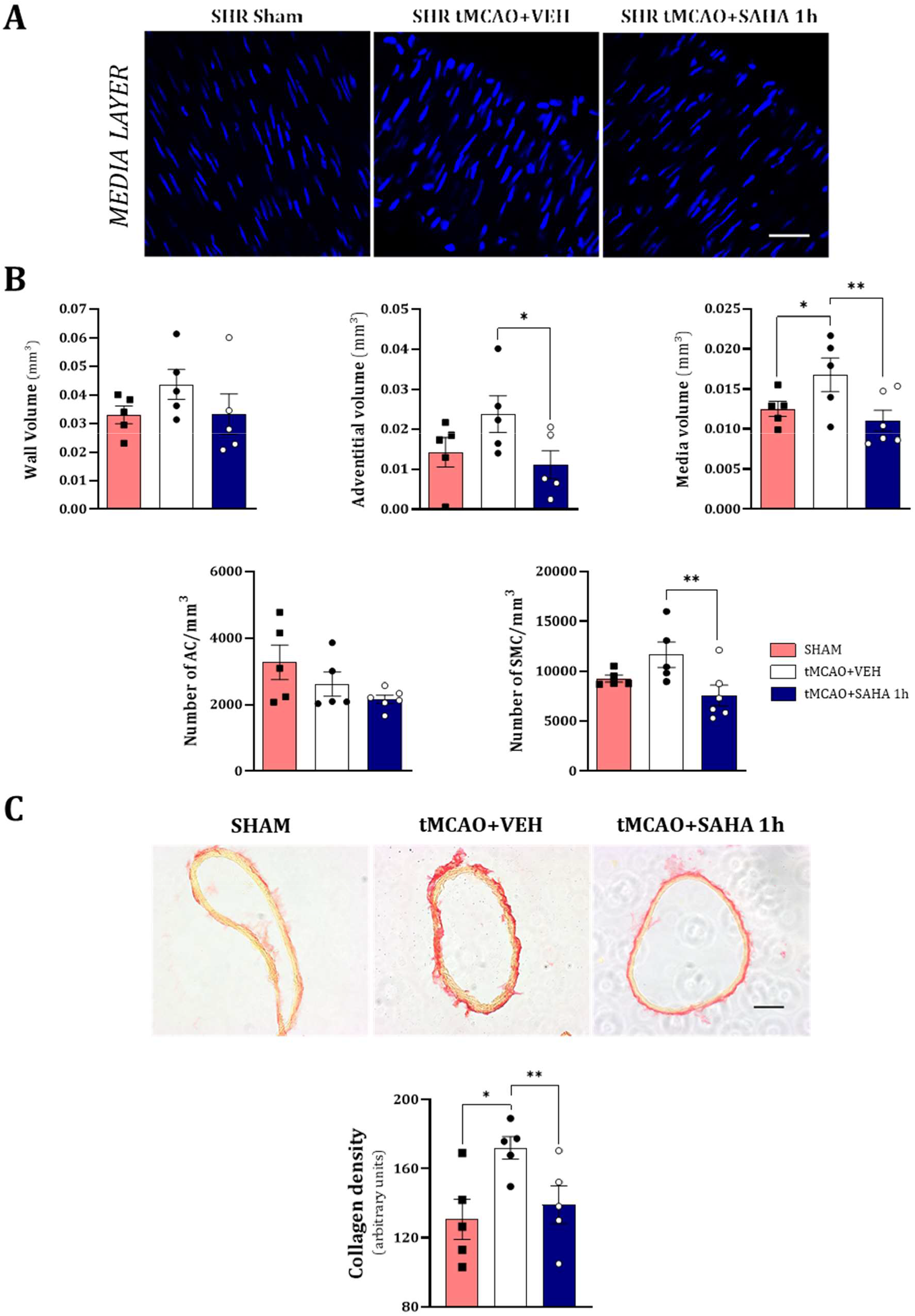
Influence of cerebral ischaemia (90 min)/reperfusion (24 h) (tMCAO) and SAHA treatment at 1 h of reperfusion on the cellular distribution within the MRA wall from hypertensive rats. The arteries were incubated with Hoechst 33342 (10 μg/ml) to stain cell nuclei (blue), and stacks of serial optical slices were taken with a laser scanning confocal microscope. (A) Representative media layer-level images obtained from confocal Z-stacks of pressurized intact segments of MRA are shown. (B) Comparison of morphological parameters. AC, adventitial cells; SMC, smooth muscle cells. Results are the mean ± SEM from SHAM (*n* = 5), tMCAO + VEH (*n* = 5), and tMCAO + SAHA 1h (*n* = 6). \**p* < 0.05; \*\**p* < 0.01 by one-way ANOVA with Bonferroni’s *post hoc* test. (C) Representative photomicrographs (top) and quantitative analysis (bottom) of total MRA collagen staining using Picrosirius Red. Scale bar 50 μm. Results are the mean ± SEM from SHAM (*n* = 5), tMCAO + VEH (*n* = 5), and tMCAO + SAHA 1h (*n* = 6). \**p* < 0.05; \*\**p* < 0.01 by one-way ANOVA with Bonferroni’s *post hoc* test.

To investigate whether alterations in arterial wall collagen played a role in cerebral I/R-induced MRA wall hypertrophy and the protective effects of SAHA, we quantified total collagen content by picrosirius red staining (Figure 5C). Notably, cerebral I/R significantly increased collagen deposition in the vessel walls, a change prevented by SAHA treatment. Overall, these findings suggest that the cerebral I/R-induced increase in media volume leading to MRA hypertrophy involves collagen deposition, which is mitigated by SAHA treatment.

### 3.4. Cerebral I/R induces an increase in both circulating and MRA oxidative stress levels that are reduced by SAHA treatment

To identify potential determinants of peripheral MRA remodelling, we measured plasma 2-EOH levels by HPLC, as an indirect indicator of circulating O_2_^·−^levels (Figure 6A), and DHE-derived fluorescence along the MRA wall to evaluate *in situ* oxidative stress (Figure 6B). Cerebral I/R induced an increase in plasma 2-EOH levels, an effect that was prevented by SAHA treatment, particularly when administered at 4 h (*p* = 0.06) and 6 h (*p* < 0.05) after reperfusion onset (Figure 6A). Consistently, cerebral I/R slightly increased DHE-derived fluorescence along the MRA wall, though this increase was not statistically significant (Figure 6B). However, SAHA treatment reduced DHE-derived fluorescence, regardless of the timing of administration.

**Figure 6.**
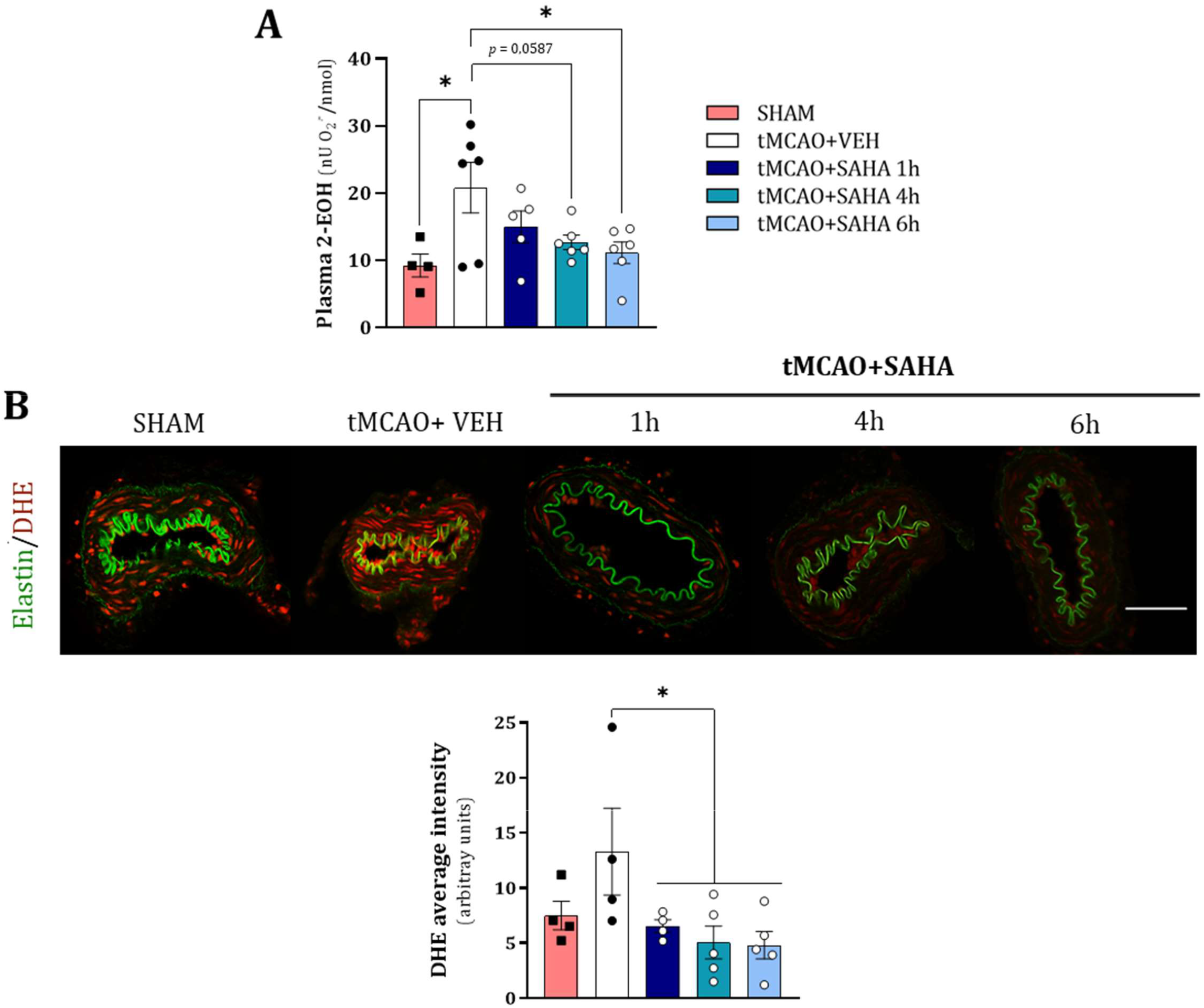
Influence of cerebral ischaemia (90 min)/reperfusion (24 h) (tMCAO) and SAHA treatment at 1, 4, or 6 h of reperfusion on circulating and MRA wall oxidative stress in hypertensive rats. (A) Analysis of 2-EOH levels in plasma by HPLC. Results are the mean ± SEM from SHAM (*n* = 4), tMCAO + VEH (*n* = 6), tMCAO + SAHA 1h (*n* = 5), tMCAO + SAHA 4h (*n* = 6), and tMCAO + SAHA 6h (*n* = 6). \**p* < 0.05 by one-way ANOVA with Bonferroni’s *post hoc* test. (B) Representative photomicrographs (top) and quantification of fluorescence intensity (bottom) of confocal microscopic MRA sections labelled with the oxidative dye dihydroethidium (DHE), which produces a red fluorescence when oxidized to ethidium bromide. Natural autofluorescence of elastin (green) is also shown. Scale bar 50 μm. Results are the mean ± SEM from SHAM (*n* = 4), tMCAO + VEH (*n* = 4), tMCAO + SAHA 1h (*n* = 4), tMCAO + SAHA 4h (*n* = 5), and tMCAO + SAHA 6h (*n* = 5). \**p* < 0.05 by one-way ANOVA with Bonferroni’s *post hoc* test.

### 3.5. SAHA treatment exerts long-term cerebroprotection

To determine whether MRA alterations are reversible or sustained over time, and if early SAHA treatment during reperfusion provides long-term cerebroprotection and beneficial effects on MRAs, SHRs underwent 90 min of MCA occlusion followed by 8 days of reperfusion. A single dose of SAHA (50 mg/kg) or an equivalent volume of vehicle was administered 4 h after the onset of reperfusion. Both body weight and SBP were similar between the two experimental conditions before the surgery, as well as on day 1 and day 8 after reperfusion (Table 1). A decrease in body weight and SBP was observed in both groups at 1 day after reperfusion onset when compared to pre-operative values, but only reached significant differences in the case of SBP. By day 8, SBP values had recovered to the levels before surgery in both groups, while body weight only recovered in the group treated with vehicle. During ischaemia (90 min) and the initial recovery at reperfusion (15 min), cCBF values were comparable between the two experimental conditions. In addition, the incidence of hyperaemia (defined as a ≥ 20% increase in cCBF compared to previous basal levels) during reperfusion, a factor associated with worse outcomes after tMCAO [38], was similar in the two groups (Table 1).

**Table 1.**
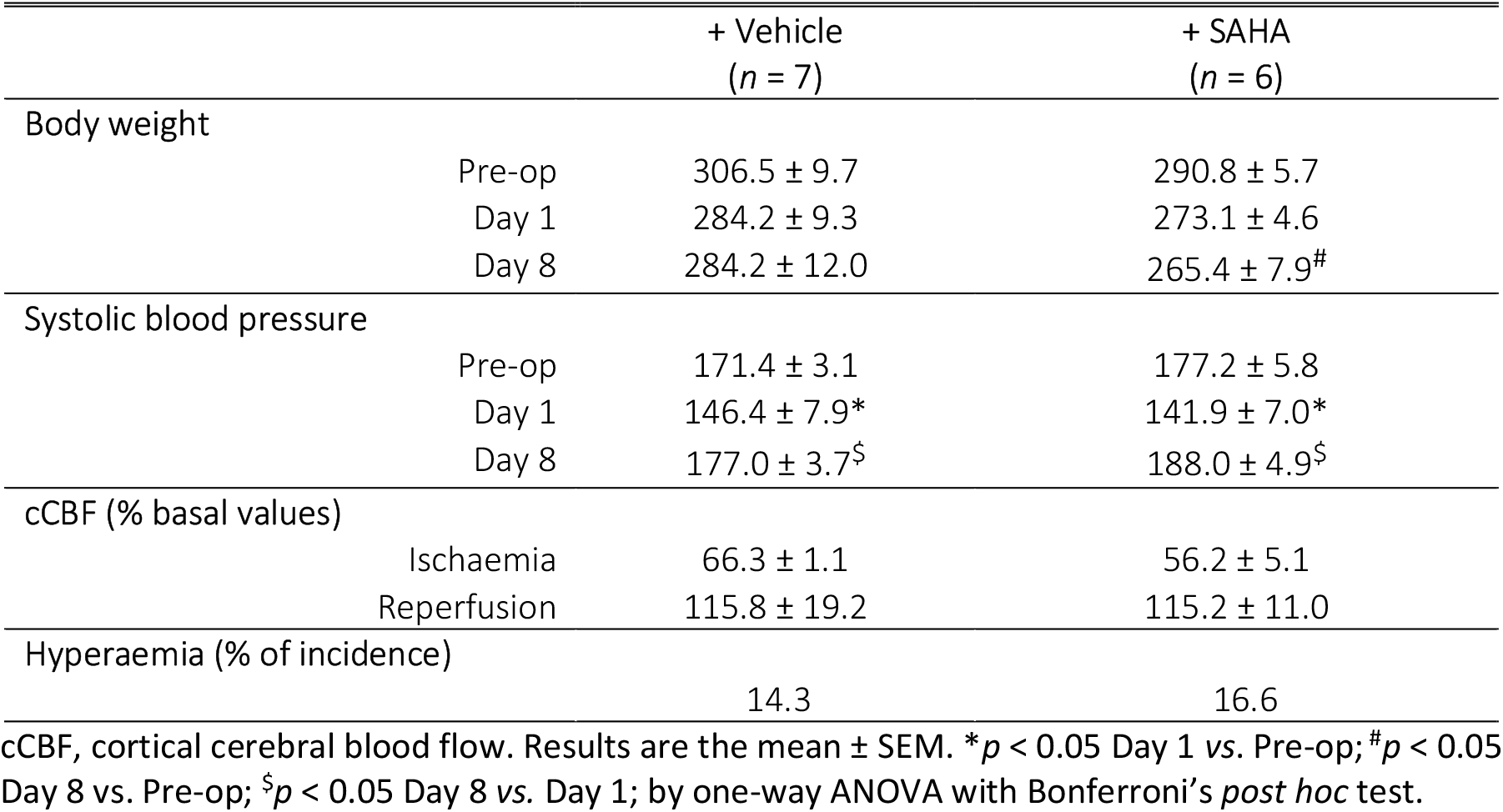
Changes in physiological parameters before (Pre-op) and after (1-8 days) the operation, and cortical cerebral blood flow (cCBF) percentage during cerebral ischaemia (90 min) and reperfusion (first 15 min), relative to basal levels.

|  |  | + Vehicle<br>(n = 7) | + SAHA<br>(n = 6) |
| --- | --- | --- | --- |
| **Body weight** |  |  |  |
| | Pre-op | 306.5 ± 9.7 | 290.8 ± 5.7 |
| | Day 1 | 284.2 ± 9.3 | 273.1 ± 4.6 |
| | Day 8 | 284.2 ± 12.0 | 265.4 ± 7.9<sup>#</sup> |
| **Systolic blood pressure** |  |  |  |
| | Pre-op | 171.4 ± 3.1 | 177.2 ± 5.8 |
| | Day 1 | 146.4 ± 7.9* | 141.9 ± 7.0* |
| | Day 8 | 177.0 ± 3.7<sup>§</sup> | 188.0 ± 4.9<sup>§</sup> |
| **cCBF (% basal values)** |  |  |  |
| | Ischaemia | 66.3 ± 1.1 | 56.2 ± 5.1 |
| | Reperfusion | 115.8 ± 19.2 | 115.2 ± 11.0 |
| **Hyperaemia (% of incidence)** |  |  |  |
|  |  | 14.3 | 16.6 |

cCBF, cortical cerebral blood flow. Results are the mean ± SEM. \* $p < 0.05$ Day 1 vs. Pre-op; <sup>#</sup> $p < 0.05$ Day 8 vs. Pre-op; <sup>§</sup> $p < 0.05$ Day 8 vs. Day 1; by one-way ANOVA with Bonferroni's *post hoc* test.

The potential of SAHA to mitigate long-term I/R brain injury following tMCAO was evaluated longitudinally using MRI scans obtained at 1- and 8-days post-injury (Figure 7A). Quantitative analysis of high-resolution T_2_w images revealed that infarct volumes in both groups spontaneously decreased from day 1 to day 8, particularly in the subcortical area (Figure 7A and 7B). A single dose of SAHA significantly reduced infarct volume on both days 1 and 8 after reperfusion onset (Figure 7A and 7B). Specifically, cortical damage was diminished by SAHA treatment both at 1 and 8 days after the injury compared to vehicle-treated animals, whereas only a trend (*p* = 0.068) was observed for subcortical damage at the early phase. Similarly, SAHA treatment prevented cortical oedema formation one day after the ischaemic insult, but this effect was not observed in the subcortical region (Figure 7C). Cortical and subcortical oedema decreased in both experimental groups 8 days after the injury (Figure 7C). Moreover, SAHA treatment significantly reduced the 9-point neurological score 8 days after the onset of reperfusion, while a slight, non-significant reduction was observed on day 1 (Figure 7D). Despite this, similar to findings observed at 1-day post-stroke in a prior study [27], SAHA did not prevent the progressive impairment in motor coordination induced by cerebral I/R (Figure 7E).

**Figure 7.**
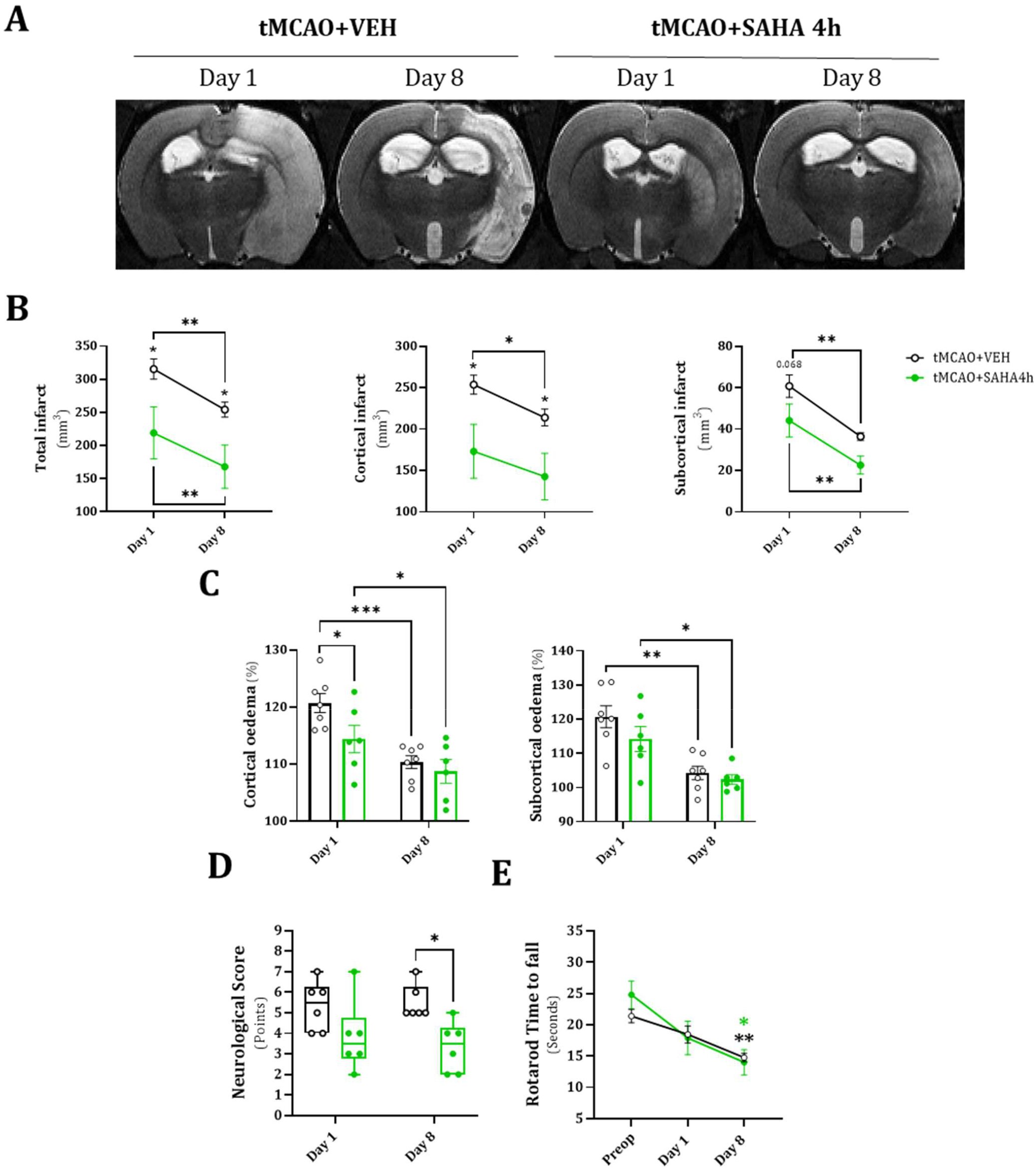
Beneficial effect of SAHA on brain damage after cerebral ischaemia (90 min)/reperfusion (1 and 8 days) (tMCAO) in hypertensive rats. Animals received a single dose of SAHA (50 mg/kg; i.p.) or vehicle at 4 h after the onset of reperfusion. (A) Representative *in vivo* high resolution T_2_-weighted MRI obtained at 1 and 8 days after reperfusion onset. (B) Quantitative analysis of total, cortical, and subcortical infarct volumes. (C) Extent of cortical and subcortical oedemas. (D) Neurological score and (E) Rotarod performance. Results are the mean ± SEM (B, C, E) or median [Q1; Q3] (D) from tMCAO + VEH (*n* = 7), and tMCAO + SAHA 4 h (*n* = 6). \**p* < 0.05; \*\**p* < 0.01; \*\*\**p* < 0.001 by repeated (B, E) or regular (C, D) measures two-way ANOVA with Tukey’s *post hoc* test.

### 3.6. Long-term persistence of cerebral I/R-induced functional and structural alterations in MRAs: minor influence of SAHA treatment

To evaluate the impact of cerebral ischaemia followed by 8 days of reperfusion and *in vivo* SAHA administration on the properties of MRA from SHR, we first assessed arterial function. In all experimental conditions, there were no notable differences in KCl (100 mM)-induced constrictions (results not shown). The impairment in U46619 contractions observed at 1 day of reperfusion was maintained after 8 days and was partially prevented by SAHA treatment (Figure 8A). Relaxations to ACh remained similar across all experimental groups, despite a slight increase in relaxation observed at a low (10^−8^ M) concentration in the group treated with SAHA (Figure 8B).

**Figure 8.**
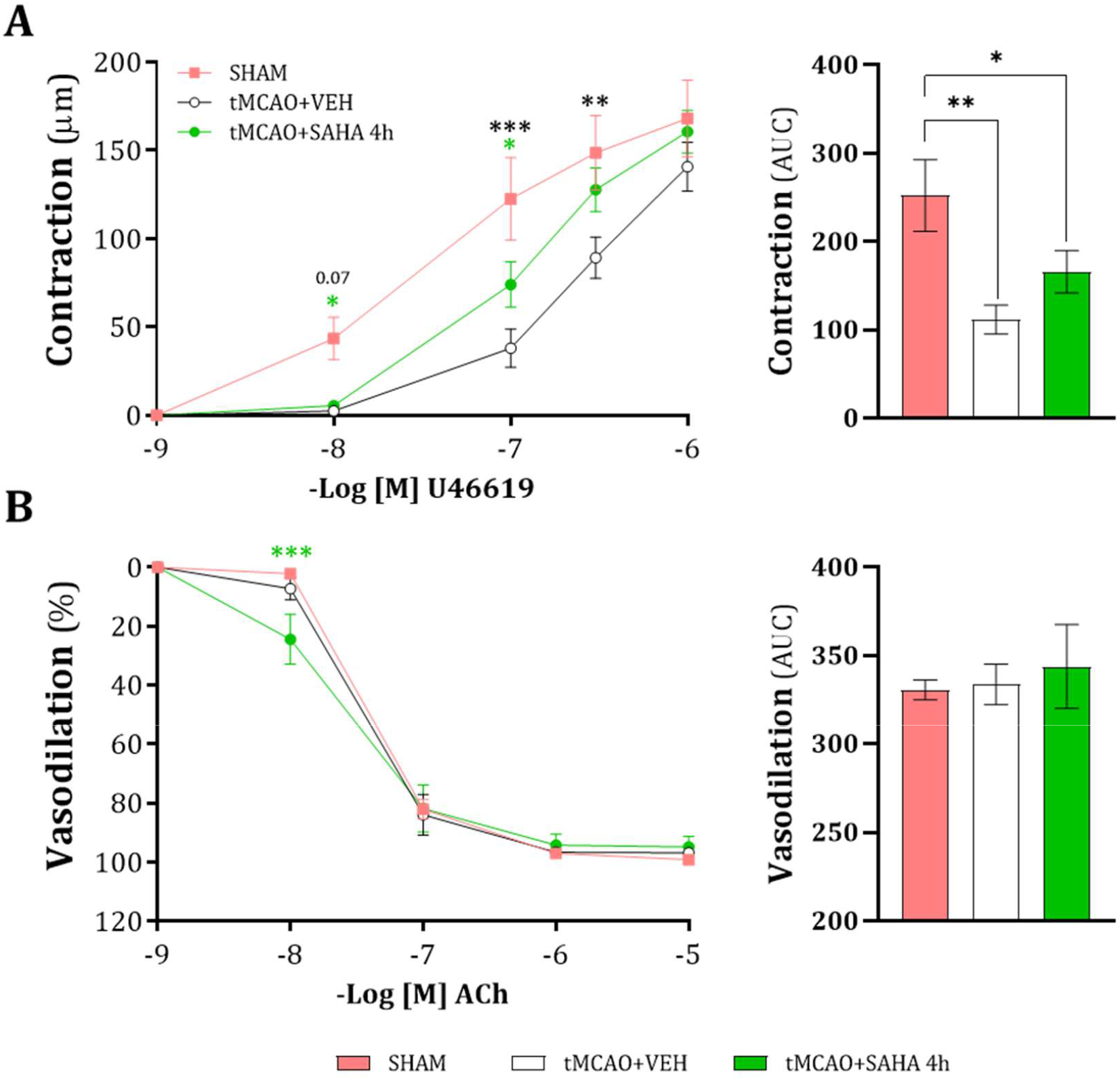
Influence of cerebral ischaemia (90 min)/reperfusion (8 days) (tMCAO) and SAHA treatment on MRA concentration-response curves to (A) U46619 (ACh) and (B) acetylcholine (ACh) in hypertensive rats. The analysis of area under the curve (AUC) is presented on the right. Results are the mean ± SEM from, SHAM (*n* = 5), tMCAO + VEH (*n* = 4), tMCAO + SAHA 1h (*n* = 6). \**p* < 0.05; \*\**p* < 0.01; \*\*\**p* < 0.001 by repeated measures two-way ANOVA with Tukey’s *post hoc* test (left) or one-way ANOVA with Bonferroni’s *post hoc* test (right).

On the other hand, arterial diameters under passive conditions (0 Ca^2+^-KH solution) were unaffected by both cerebral I/R and SAHA treatment (results not shown). Notably, cerebral I/R resulted in an increase in MRA wall thickness (Figure 9A), at intraluminal pressures ranging from 3 to 40 mmHg, and cross-sectional area (Figure 9B) compared to sham-operated rats, a phenomenon unaffected by SAHA treatment. In contrast, the wall-to-lumen ratio remained unchanged (Figure 9C). Taken together, after 8 days of reperfusion, MRAs show non-reversible impaired contractile responses to the thromboxane A2 receptor agonist U46619, and SAHA has minimal impact on these chronic contractile impairments. Additionally, hypertrophic remodelling of MRAs persists long-term, with no discernible effect from SAHA treatment.

**Figure 9.**
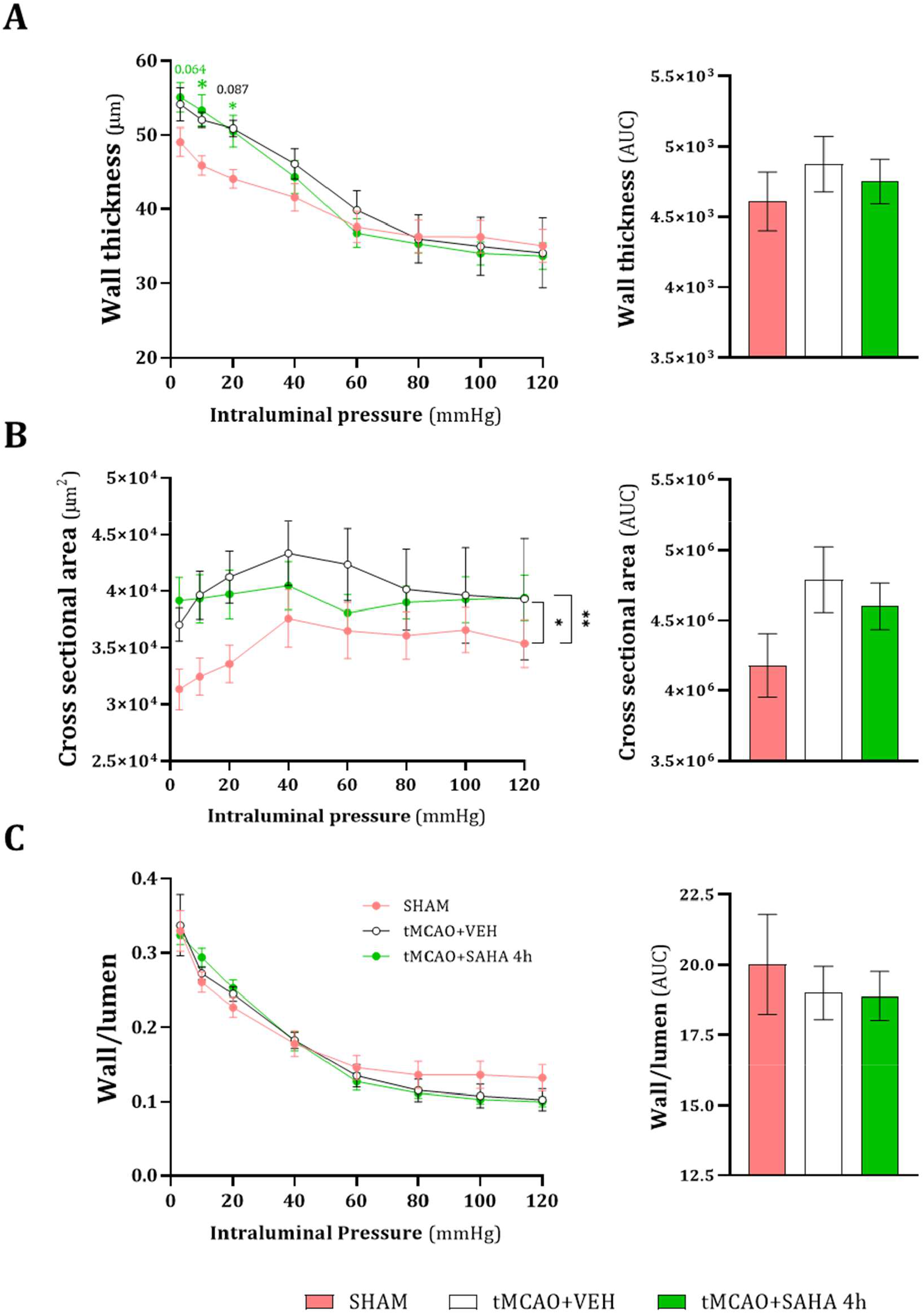
Influence of cerebral ischaemia (90 min)/reperfusion (8 days) (tMCAO) and SAHA treatment at 4 h of reperfusion on MRA structural properties obtained in passive conditions (0 Ca^2+^ – Krebs-Henseleit solution) in hypertensive rats. (A) Wall thickness – intraluminal pressure. (B) Cross-sectional area – intraluminal pressure. (C) Wall/lumen ratio – intraluminal pressure. The analysis of area under the curve (AUC) is presented on the right. Results are the mean ± SEM from SHAM (*n* = 5), tMCAO + VEH (*n* = 4), and tMCAO + SAHA 4h (*n* = 6). \**p* < 0.05; \*\**p* < 0.01 by repeated measures two-way ANOVA with Tukey’s *post hoc* test.

## 4. Discussion

Cerebral I/R injury primarily affects the brain but often leads to secondary damage in peripheral organs, worsening stroke outcomes and elevating mortality rates [5]. Understanding the mechanisms of cerebral I/R-induced peripheral injury and identifying effective treatments is crucial for improving outcomes, especially in hypertensive patients in whom post-stroke damage is aggravated. This study investigated the effects of cerebral I/R on MRAs from hypertensive rats and assessed the protective long-term effects of a single dose of SAHA administered during reperfusion. Following cerebral I (90 min)/R (24 h), MRAs from SHR exhibited impaired contractile responses to the thromboxane A2 receptor agonist U46619, persisting even after 8 days. Arterial structure analysis revealed stroke-induced MRA hypertrophy likely due to increased collagen deposition. Notably, SAHA treatment did not improve U46619 contractions but mitigated oxidative stress and collagen accumulation, thus preventing MRA remodelling at 24 h of reperfusion. Furthermore, treatment with SAHA induced cerebroprotective effects that extended long-term. In contrast, SAHA had minimal impact on chronic MRA contractile impairments and vascular remodelling. Taken together, these findings suggest that cerebral I/R causes sustained peripheral artery changes in hypertension. While SAHA treatment offers protection against brain injury and early vascular damage, it cannot fully restore long-term peripheral artery dysfunction.

After a transient ischaemic insult, functional and structural alterations occur in rat MCAs [39, 34, 31]. However, the impact of a transient cerebral ischaemic event on peripheral arteries is less understood, with as far as we know only one study reporting endothelial dysfunction in MRAs from normotensive rats [23]. In addition, there is no evidence on whether cerebral I/R exacerbates peripheral vascular changes induced by hypertension. In the present study, cerebral I/R did not modify endothelial function and KCl-induced contractions but permanently impaired U46619-induced contractions, suggesting a sustained disturbance in thromboxane A2-mediated signalling over time. Given the deleterious roles of thromboxane signalling in stroke, an impairment of thromboxane A2 vascular responses may be an adaptive response to counterbalance potentially excessive thromboxane A2-derived signalling [20]. After cerebral I/R, there is an increase in thromboxane receptors in ipsilateral brain tissue, likely via microglia activation [19]. Despite this, similar impairments to those observed in the present study in U46619-induced contractions were previously noted in ovariectomized rat pial arteries after transient global cerebral ischaemia [40]. It is worth mentioning that MRAs from old normotensive Wistar-Kyoto rats and old SHR show decreased reactivity to U46619 due to reduced expression of thromboxane receptors and increased circulating levels of the endogenous thromboxane receptor antagonist 8,9-epoxyeicosatrienoic acid [41]. A similar process may occur in MRAs after cerebral I/R. Nevertheless, further studies should confirm this hypothesis and investigate the specific mechanisms by which cerebral I/R diminishes thromboxane A2-mediated contractions in these arteries.

For the first time, we report that cerebral I/R induces increases in cross-sectional area, wall thickness, and wall-to-lumen ratio in MRAs from SHR. These findings partly support previous research showing exacerbated hypertrophic remodelling in MCAs from SHR following cerebral I/R [31]. Notably, the observed hypertrophic remodelling in MRAs following stroke underscores significant pathological implications. These alterations can contribute to long-term cardiovascular events by increasing vascular resistance, impairing vascular function, and potentially promoting atherosclerosis and thrombosis [42, 43]. Therefore, mitigating these changes is critical for preventing cardiovascular complications in hypertensive patients following cerebral ischaemic events. To investigate the cellular basis of these changes, we found that MRA hypertrophic remodelling was primarily due to an increase in the media layer volume without a corresponding increase in SMC proliferation, although a tendency was observed. This led us to examine whether changes in collagen content could be involved. Consistently, we detected increased collagen deposition in MRAs after cerebral I/R, indicating that this extracellular matrix protein is likely the main contributor to the observed vascular remodelling. Importantly, we have demonstrated that these alterations are not entirely reversible, as evidenced by the persistent MRA hypertrophy observed 8 days post-injury.

Histone deacetylase inhibitors are gaining attention as a promising therapeutic avenue due to their ability to modulate the expression of neuroprotective and regenerative genes [44]. Specifically, SAHA, a pan-specific HDAC inhibitor approved for the treatment of cutaneous lymphoma, has demonstrated cerebroprotective effects in models of both permanent [24, 25] and transient [26, 27] cerebral ischaemia. Notably, some studies have described the anti-inflammatory [45, 46] and antioxidant [47] properties of SAHA in peripheral systems, as well as its ability to attenuate SMC proliferation in an *in vitro* model [48]. Here, we demonstrate the potential of SAHA in preventing cerebral I/R-induced MRA hypertrophic remodelling in hypertensive rats. After 24 h of reperfusion, SAHA treatment effectively halted wall hypertrophy and attenuated increases in the wall-to-lumen ratio when administered within 4 h of the onset of reperfusion. Our findings suggest that this protective effect involves both a reduction in the number of SMCs and a decrease in collagen content. This aligns with previous research indicating that SAHA, in addition to other HDAC inhibitors [49, 50], reduce SMC proliferation (for review, see [51]), and that HDAC inhibitors (e.g., Trichostatin A) decrease collagen production in isolated cardiac fibroblasts [52].

Reactive oxygen species play a crucial role in maintaining vascular homeostasis under physiological conditions [53]. However, under pathological situations, ROS contribute to multiple damaging processes, namely inflammation, hypertrophy, proliferation, apoptosis, and migration, all of which perpetuate hypertension-induced vascular damage [54]. Our results suggest that administering SAHA during reperfusion exerts antioxidant actions by reducing O_2.-_ levels in plasma and decreasing *in situ* oxidative stress in MRAs. These findings may explain the observed decrease in collagen content induced by SAHA treatment. Proinflammatory genes regulated by ROS, mediated by redox-sensitive transcription factors, contribute to vascular remodelling at least partly through SMC growth, proliferation, and collagen accumulation [55]. Antioxidant treatments have shown promise in counteracting SMC alterations and reducing collagen accumulation in inflammatory contexts. Therefore, by targeting oxidative stress, SAHA administration during reperfusion may mitigate collagen accumulation, thereby buffering short-term MRA remodelling.

The short-term cerebroprotective effects of SAHA have been previously demonstrated in various murine models of ischaemic stroke [24, 26, 27]. However, only one study has reported potential long-term neuroprotective effects of this drug in a normotensive mouse model of permanent occlusion [25]. In the present study, assessing brain infarct evolution by MRI, we confirmed the previously reported cerebroprotective effects of SAHA within 24 h of reperfusion in hypertensive rats [27]. Notably, our longitudinal examination is the first to demonstrate the sustained efficacy of a single SAHA administration at 4 h of reperfusion in hypertensive animals, extending its protective effects up to 8 days post-stroke. Within our experimental framework, SAHA demonstrated neuroprotective benefits, including improvements in neurological function and significant reductions in both infarction size and oedema formation. In summary, our findings suggest that SAHA shows promise as an effective therapeutic agent for managing brain damage after stroke. However, the present study also aimed to investigate whether SAHA could mitigate cerebral I/R-induced peripheral vascular changes. While SAHA administration partially improved thromboxane A2-induced contractions 8 days post-cerebral I/R, it did not yield significant benefits for vascular remodelling, as MRA hypertrophy did not differ from that in the vehicle group. These results imply that the complete prevention of MRA remodelling seen 1-day post-stroke with SAHA treatment reverts over time. Moreover, the treatment did not normalize body weight, confirming incomplete recovery. These findings suggest that while SAHA provides long-lasting cerebroprotection, it may not completely prevent peripheral dysfunction in the long term. We hypothesize that a single dose administered during reperfusion might not be sufficient and that additional repetitive doses may be required to mitigate peripheral damage.

Some limitations should be acknowledged. This study only utilized male young rats, leaving uncertainty about whether the observed effects on MRAs are representative of responses in females or in older animals. Furthermore, the possibility exists that other blood vessels/organs could experience either no changes or different changes compared to MRAs. In addition, the focus of the study was specifically on the effects of cerebral I/R injury in hypertensive SHR, which raises questions about the generalizability of the findings to other models of hypertension or to individuals with different comorbidities (e.g., diabetes, obesity…). Further research involving diverse animal models, considering both genders, older ages, and different comorbidities, is necessary to comprehensively understand the mechanisms underlying cerebral I/R-induced peripheral injury. Such studies will help in developing effective therapeutic strategies that can address both cerebral and peripheral complications of ischaemic stroke across diverse populations.

In summary, cerebral I/R induces thromboxane A2-induced contractile dysfunction as well as hypertrophic remodelling in MRAs from SHR. SAHA administration during early reperfusion provides significant protective effects by reducing oxidative stress and collagen-induced hypertrophic remodelling after 24 h of reperfusion. However, it does not alleviate altered thromboxane A2 contractions. In addition, while SAHA offers both short- and long-term cerebroprotection, it does not fully prevent peripheral artery disturbances over time. These findings highlight the importance of monitoring peripheral artery damage following cerebral I/R and highlight the crucial role of early SAHA intervention in mitigating brain damage and certain peripheral vascular complications. However, cerebroprotective treatments may not sufficiently improve peripheral damage after a stroke over time, highlighting the need for further research to address long-term vascular health throughout the body.

## Ethics approval and informed consent

All the experiments were carried out in accordance with the guidelines established by the Spanish legislation on protection of animals used for scientific purposes (RD 53/2013) and the European Union Directive (2010/63/UE). The protocols received approval from the ethics committee of the Universitat Autònoma de Barcelona (approval code: CEAAH 4275M4).

## CRediT authorship contribution statement

**Andrea Díaz-Pérez**: Writing – review & editing, Writing – original draft, Visualization, Methodology, Investigation, Formal analysis, Data curation. **Silvia Lope-Piedrafita:** Writing – review & editing, Visualization, Methodology, Investigation, Formal analysis, Data curation. **Belén Pérez:** Writing – review & editing, Visualization, Methodology, Investigation, Formal analysis, Data curation. **Paula Vázquez-Sufuentes:** Writing – review & editing, Visualization, Methodology, Investigation. **Maria Rodriguez-Garcia:** Writing – review & editing, Visualization, Methodology, Investigation. **Ana M Briones:** Writing – review & editing, Formal analysis, Data curation. **Xavier Navarro:** Writing – review & editing, Resources, Project administration, Funding acquisition. **Clara Penas:** Writing – review & editing, Visualization, Resources, Project administration, Methodology, Investigation, Funding acquisition, Formal analysis, Data curation. **Francesc Jiménez-Altayó:** Writing – review & editing, Writing – original draft, Visualization, Validation, Supervision, Software, Resources, Project administration, Methodology, Investigation, Funding acquisition, Formal analysis, Data curation, Conceptualization.

## Acknowledgments

We are grateful to the confocal microscopy core from the Universitat Autònoma de Barcelona.

## Data availability

Data will be made available from the corresponding author upon request.

